# Strata use in a canopy-beetle community of a lowland Neotropical rainforest in southern Venezuela

**DOI:** 10.1101/2024.01.07.574520

**Authors:** Susan Kirmse

## Abstract

The stratification in tropical rainforests is well pronounced. This strongly impacts the distribution of arthropods. As part of a larger beetle survey in the northern part of the Amazonian rainforest, I analyze the characteristics of beetle species shared between the canopy and the understory. Linking the strata use of adult beetles in the complex structure of tropical rainforests with their ecological characteristics may uncover causes of the differences in diversity between vertical layers. Seventy out of a total of 862 beetle species in 45 families sampled on 23 canopy-tree species were collected also in the understory. Beetle families represented with most species in the canopy and ground samples comprise Curculionidae, Chrysomelidae, and Carabidae. In Elateridae and Scarabaeidae, the proportion of shared species between both strata amounted to ≥ 20%. In contrast, the species-rich families (≥ 20 canopy species) Cerambycidae, Mordellidae, and Buprestidae did not comprise species sampled in both strata. Adult feeding requirements and larval substrates are discussed as main reasons for the differences in strata use between these beetle families. In addition, different climatic conditions between the exposed canopy and the more uniform understory might cause a migration of adult beetles between both strata and thus, result in intermixing of beetle species from different strata within the forest.

## Introduction

The distribution of arthropods in forests is not uniform. Forest canopies are unique zones of biodiversity (Nakamura et al. 2017). Particularly, tropical rainforest canopies are famous for their richness in beetles and other arthropods (Erwin 1988, 2013; Basset et al. 2015; Nakamura et al. 2017). Analysis of Sulawesi beetle data revealed a 50% increase in species count for every additional 10 m of vertical height (Davis & Sutton 2012). Nocturnal beetles in Sulawesi showed greatest abundance, species richness, and diversity in the canopy (Davis et al. 2011). Kato et al. (1995) showed that beetle numbers increased significantly from floor to canopy and Hemiptera were more abundant in the canopy than at lower strata in a tropical rainforest in Borneo.

Typically, arthropod abundance and diversity in the upper canopy are between two and four times as great as those in the understory (Basset 2001). However, some studies found for several insect groups including Coleoptera, Formicidae, Hymenoptera, and Lepidoptera that the ground may be as speciose as the canopy (Brühl et al. 1998; Schulze et al. 2001; Stork & Grimbacher 2006; Ashton et al. 2016; Somavilla et al. 2019). In a Central Amazonian forest, Coleoptera, Diptera, and Hymenoptera had their greatest abundance at the ground level, whereas Hemiptera and Lepidoptera were more abundant in the upper levels of the canopy (Amorim et al. 2022).

Niche partitioning along vertical and elevational gradients is well known as an important mechanism in community assembly, species coexistence and consequently as diversity cue. Along a tropical elevational gradient, for instance, the niche partitioning of animals was mainly related to the functional diversity of foraging strata in plant communities (Albrecht et al. 2018). Forests are characterized by a substantial change in habitat conditions between understory and canopy strata (Shaw 2004; Basset et al. 2015; Nakamura et al. 2017). Within and between tropical lowland rainforests, the number and distinctness of vegetation strata largely vary in relation to floristic composition (Yamakura 1987; King et al. 2006). Understory tree composition varies with Neotropical and African rainforests harboring many small flowering trees, while non-reproductive juveniles of canopy trees dominate in South East Asian rainforests (LaFrankie et al. 2006). In Amazonian forests, the dominant plant species in the understory are distinct from upper canopy dominants (Laurans et al. 2014; Draper et al. 2020).

This vertical stratification is often considered to be a key factor promoting the extreme diversity of tropical forests (Oliveira & Scheffers 2019). Strata preferences may guide the occurrence of species in distinct layers. Many arthropod groups show clear patterns of stratification particularly in complex tropical rainforests (Basset et al. 2003b; Stork et al. 2015). In general, arthropod assemblages including mites (Arachnida) (Walter 1995), Collembola (Rogers & Kitching 1998), Formicidae (Longino & Nadkarni 1990; Brühl et al. 1998), Lepidoptera (DeVries et al. 1997, 2012; Schultze et al. 2001; Mena et al. 2020), or Chrysomelidae (Charles & Basset 2005) in the canopy are very distinct from those that inhabit the understory (Basset et al. 2001). Amorim et al. (2022) found in a central Amazonian tropical forest that 61.6% of all Diptera species were not sampled at the ground level indicating their stratum specificity. These patterns of stratification in tropical forests resemble that found in temperate forests (Normann et al. 2016; Weiss et al. 2019). Leksono and Nakagoshi (2015) demonstrated in a study with window traps that canopy insect communities vary among canopy and understory layer.

Roubik (1993) and Floren (2010) indicated that patterns of stratification in forests are sometimes mitigated by the preferential and seasonal use of certain food resources. However, even the same food resources in diverse strata can attract different species. Barrios (2003) compared herbivore species on saplings and trees and found only one leaf-chewing chrysomelid species occurring on both. Similarly, the composition of flower-visitor communities exhibits generally a large difference between the canopy and the understory in tropical (Roubik 1993; Nagamitsu et al. 1999; Ramalho 2004) and temperate forests (Ulyshen et al. 2010). In addition, depending on their ecology, certain guilds like scavengers occur preferential near the ground level, while many herbivores occur in the upper canopy (Basset et al. 2003b). Schleuning et al. (2011) concluded that vertical stratification could foster niche partitioning among different functional guilds. Consequently, different beetle families and guilds, respectively, should exhibit different patterns of strata use in concordance with their ecology. In this study, I analyze the strata use of canopy beetles collected on 23 Amazonian tree species and relate their strata specificity to their larval and adult feeding habitats.

## Material and methods

### Study site

The general beetle survey was conducted as part of the interdisciplinary research project “Towards an understanding of the structure and function of a Neotropical rainforest ecosystem with special reference to its canopy” organized by the Austrian Academy of Science. The study site is located in the upper Orinoco region (Venezuela, state of Amazonas) (3°10’N, 65°40’W; 105 m asl). A canopy crane was installed at the small black-water river Surumoni, a tributary of the large white-water river Orinoco. The tower crane was 42 m in height and ran on 120 m long rails. An area of about 1.4 ha was accessible with the crane’s 40 m long swing. A gondola carrying the scientists and their equipment enabled movement between the tree crowns.

The Surumoni area belongs to the Japura/Negro moist forests ecoregion (Dinerstein et al. 1995) or Imerí province (Morrone 2014) that extends from Brazil to southern Venezuela, Colombia, and Peru. The vegetation in this remote part of lowland moist forest in the northern Amazon basin is classified as terra firme (Prance 1979). The annual precipitation is about 3100 mm, although year to year fluctuations of about 500 mm occur (Anhuf et al. 1999). There is strong peak in the precipitation from May to July, then a lower peak in September and October. Due to the El Niña phenomenon in 1996, there was extremely high rainfall, whereas El Niño caused a strong dry period in the change of 1997 to 1998. The average annual temperature in the study area is ca. 26°C, usually with slight variations between the coolest month (25°C) and the warmest month (26.5°C). Maximum temperatures at day may reach 30.5°C and drop to only 20-21°C during the night.

Altogether 322 species of higher plants were identified in the 1.4 ha crane plot belonging to 208 genera from 78 families (Wesenberg 2004). The forest is frequently interrupted by light gaps due to irregular crown closure. The upper canopy ranges usually from 25 to 27 m in height. A few emergent trees rise to a height of 35 m. The Surumoni study site is average in tree species richness with an estimated number of 92 species per hectare. There were more than 800 trees ≥ 10 cm DBH (diameter at breast height) belonging to 141 tree species within the crane plot (Wesenberg 2004).

Frequent species in the tree fraction with a DBH of ≥ 10 cm were *Dialium guianense* (Aubl.) Sandwith (Fabaceae), *Goupia glabra* Aubl. (Goupiaceae), *Ocotea aff. amazonica* (Meisn.) Mez (Lauraceae), *Oenocarpus bacaba* Mart. (Arecaceae), and *Ruizterania trichanthera* (Spruce ex Warm.) Marc.-Berti (Vochysiaceae). Abundant species of the upper layer included *G. glabra*, *R. trichanthera*, and *D. guianense*. Abundant species of the middle layer were, for instance, *O. bacaba*, *Couma utilis* (Mart.) Müll. Arg. (Apocynaceae), and *Podocalyx loranthoides* Klotzsch (Picrodendraceae). The underlayer growth (5-10 m) consisted of young *G. glabra* and *O. bacaba*. Epiphytes and hemiepiphytes were rare compared to other moist forests comprising 53 species with Araceae reaching the highest abundance (Engwald et al. 2000). The herb layer was well developed. It was dominated by ferns of the families Hymenophyllaceae and Metaxyaceae as well as small palms of the genera *Geonoma* Willd. and *Bactris* Jacq. ex Scop. Other abundant plants included Rubiaceae (*Psychotria* L. spp. and *Faramea* Aubl.), Melastomataceae and Maranthaceae (*Ischnosiphon* Körn. spp.) as well as Heliconiaceae (Wesenberg 2004). The forest floor was sparsely covered with leaf litter.

### Field studies

The field study was carried out between 1997 and 1999. Observations and collections of canopy beetles comprised the following periods: September to November 1997; May to August and December 1998; January to April 1999, and thus, cover combined a full year. Additional aerial trap collection was performed in October 1999 and targeted a single tree species. The periods of sampling and observation of adult beetles in the understory vary slightly due to different canopy access: August to November 1997; April to August and December 1998; January to March 1999. License and authorization were issued under the number 15-1277 by Servicio Autonoma de Fauna, Ministerio del Ambiente y de los Recursos Naturales Renovables, Caracas, Venezuela.

The beetle survey includes 23 tree species representing 13 plant families in the upper (approximately 25–30 m height) and middle (approximately 18–25 m height) canopy (Figure 1). The trees selected for the beetle survey were either completely free from epiphytes and lianas or bore only small ones to minimize errors of beetle host associations. Parts of the tree crowns sampled comprise leaves, small twigs, flowers, and fruits. The trees chosen were searched regularly for Coleoptera during the day and the night. Observed beetles were captured by net, hand or through branch and foliage beating. These collection methods were not structured to provide quantitative data. In addition, 10 aerial traps were used to collect flying beetles. These flight interception traps consisted of two clear acrylic panels fixed in a cross with each a length of 30 cm and a height of 25 cm. A plastic tube ending in a container filled with water mixed with a surface tension-diminishing detergent collected the beetles. The trapped beetles were removed every second day.

**Figure 1.**
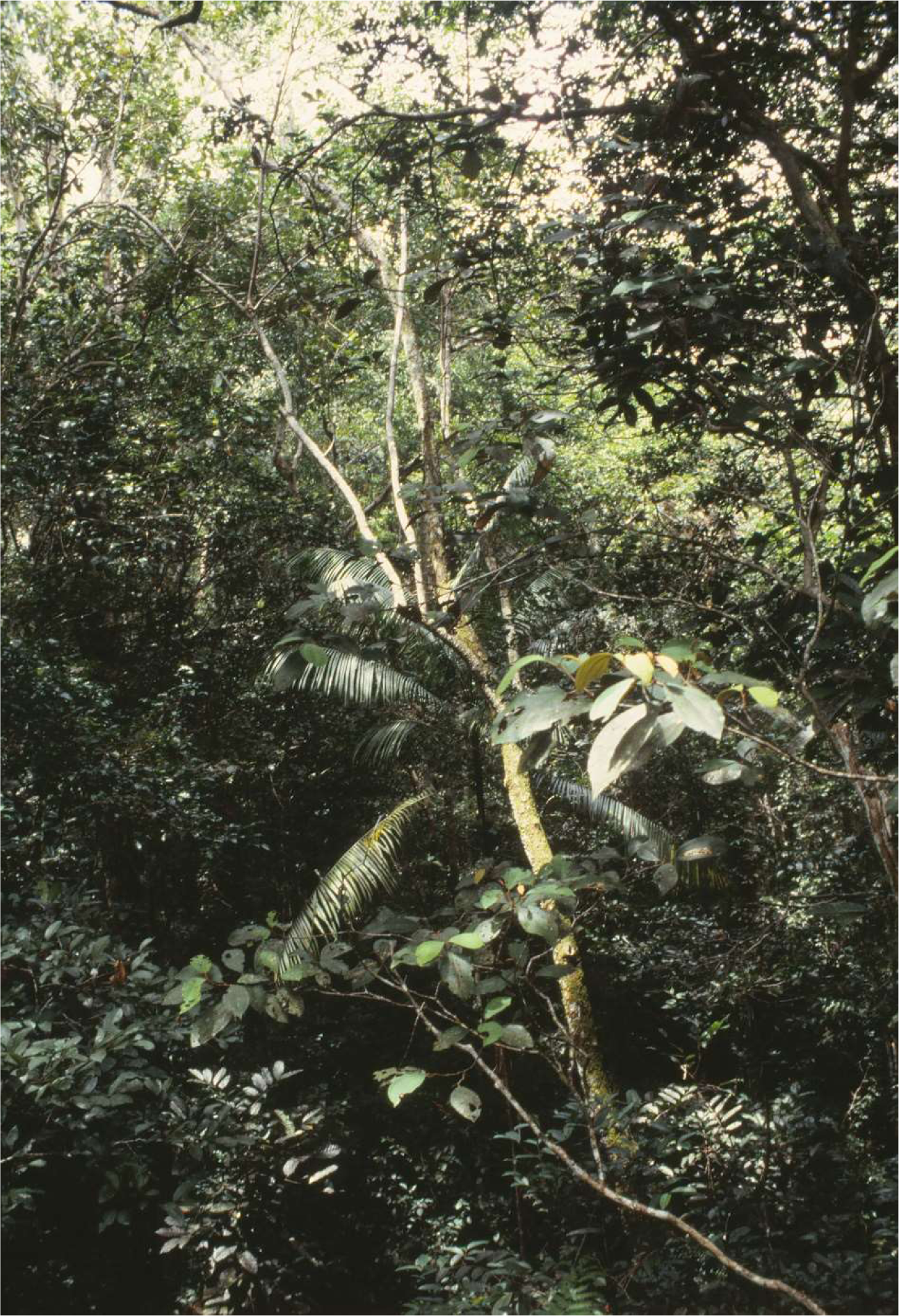
Canopy of the Surumoni crane plot in a lowland tropical rainforest in Venezuela, February 1999.

The sampling regime for understory (Figure 2) beetles included non-quantitative hand collection and pitfall trapping. Pitfall traps (each three per site; diameter: 6 cm) were filled up to one third of their height with water mixed with a surface tension-diminishing detergent. The captured beetles were extracted two or three times per week. The pitfall traps were installed below fruiting trees of *G. glabra* (all periods), *Miconia poeppigii* Triana (Melastomataceae) (December 1998 to March 1999), *O. bacaba* (May 1998), *Ficus* L. sp. (Moraceae) (September 1997), *Protium* cf. *spruceanum* (Benth.) Engl. (Burseraceae) (August to September 1997; May to August 1998), *Alibertia* cf. *latifolia* (Benth.) K. Schum. (Rubiaceae) (March 1999; July 1998), and *Esenbeckia* Kunth sp. (Rutaceae) (August to September 1997). Hand collection was conducted around the fruit falls and along a 40 m long transect on a small path every second day and night. Herbs, young trees, and trunks up to a height of about 1.5 m and the ground were checked visually for adult beetles and collected manually.

**Figure 2.**
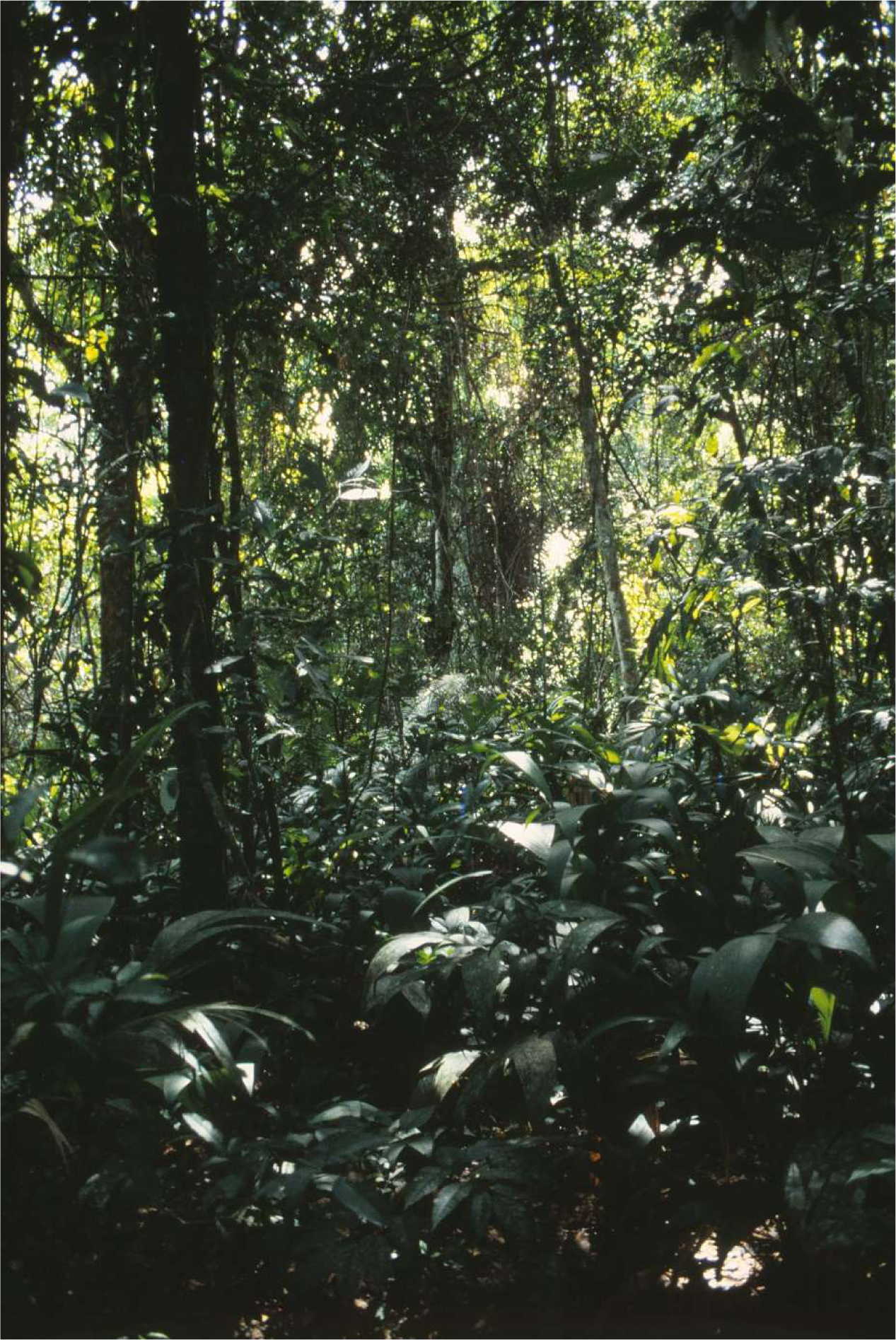
Understory in the study area of the Surumoni crane plot in a lowland tropical rainforest in Venezuela, May 1998.

### Data processing and analyses

The collected beetles were kept in 70% ethanol. The beetles were assigned to morphospecies and later partly identified by experts: Cantharidae by Alistair S. Ramsdale; Chrysomelidae in part by Shawn M. Clark, Wills Flowers, David Furth, and Lev N. Medvedev; Curculionidae by Sergio Antonio Vanin; Elateridae by Paul J. Johnson; Lampyridae by Vadim R. Viviani; Scarabaeidae by Brett C. Ratcliffe; and Tenebrionidae by Martin Lillig. The family-group taxonomic classification follows Bouchard et al. (2011). Voucher specimens of collected beetles are deposited in the Museo del Instituto de Zoología Agrícola ‘Francisco Fernandez Yepez’, Maracay, Venezuela.

I included every individual adult beetle collected on the 23 target tree species in the data set. In a first step, I added all individuals of canopy-beetle species collected on the ground/understory to this data set (Table 1). Afterwards, all unique singletons were omitted from the data set for further analyses (Table 2). In a third step, I added all congenerics collected on the ground/understory to the identified canopy genera (Table 3). While all species of Cantharidae, Carabidae, Elateridae, Lampyridae, and Scarabaeidae collected in the understory were assigned to a genus, the genus identification of Chrysomelidae and Curculionidae was carried out only in part.

**Table 1.**
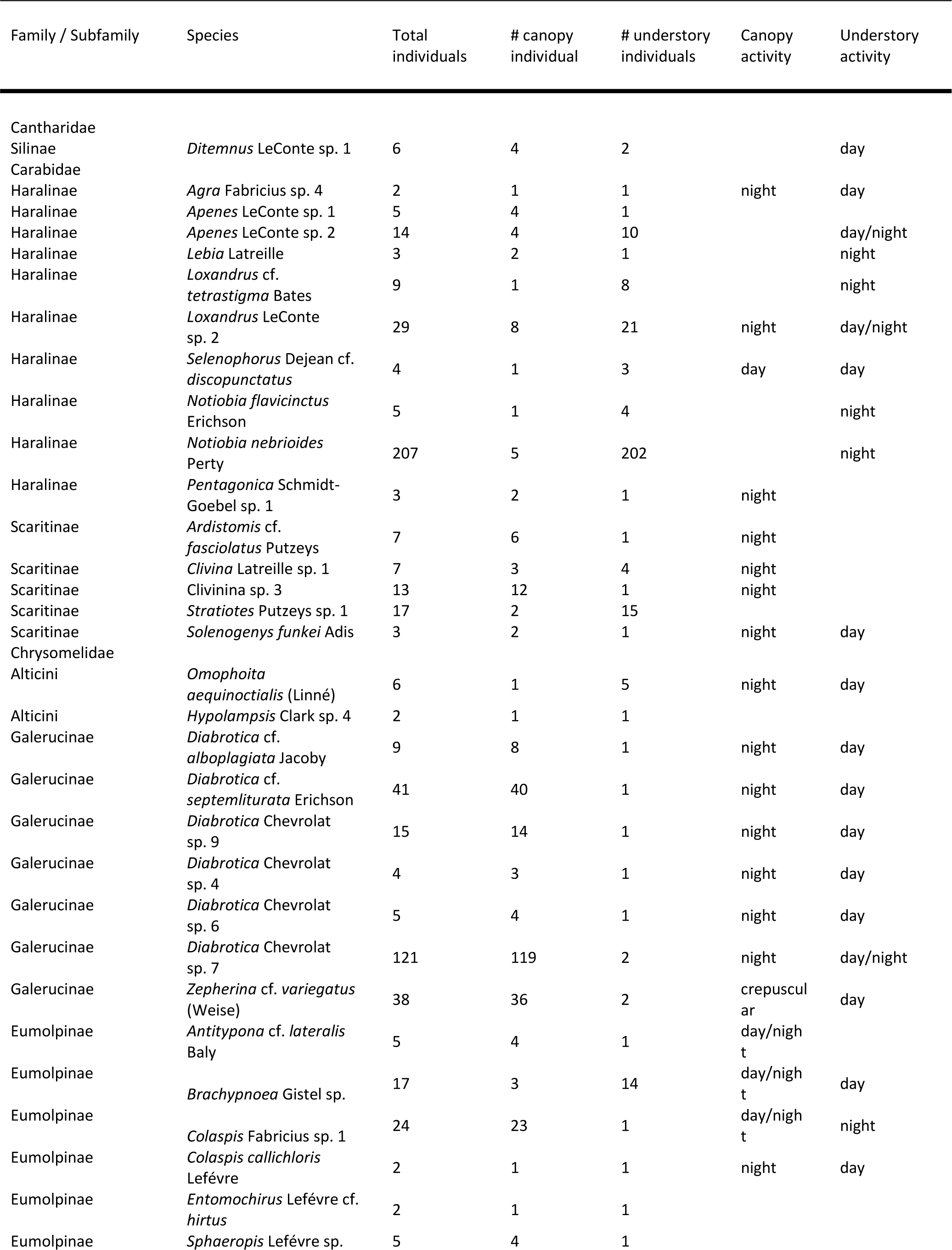
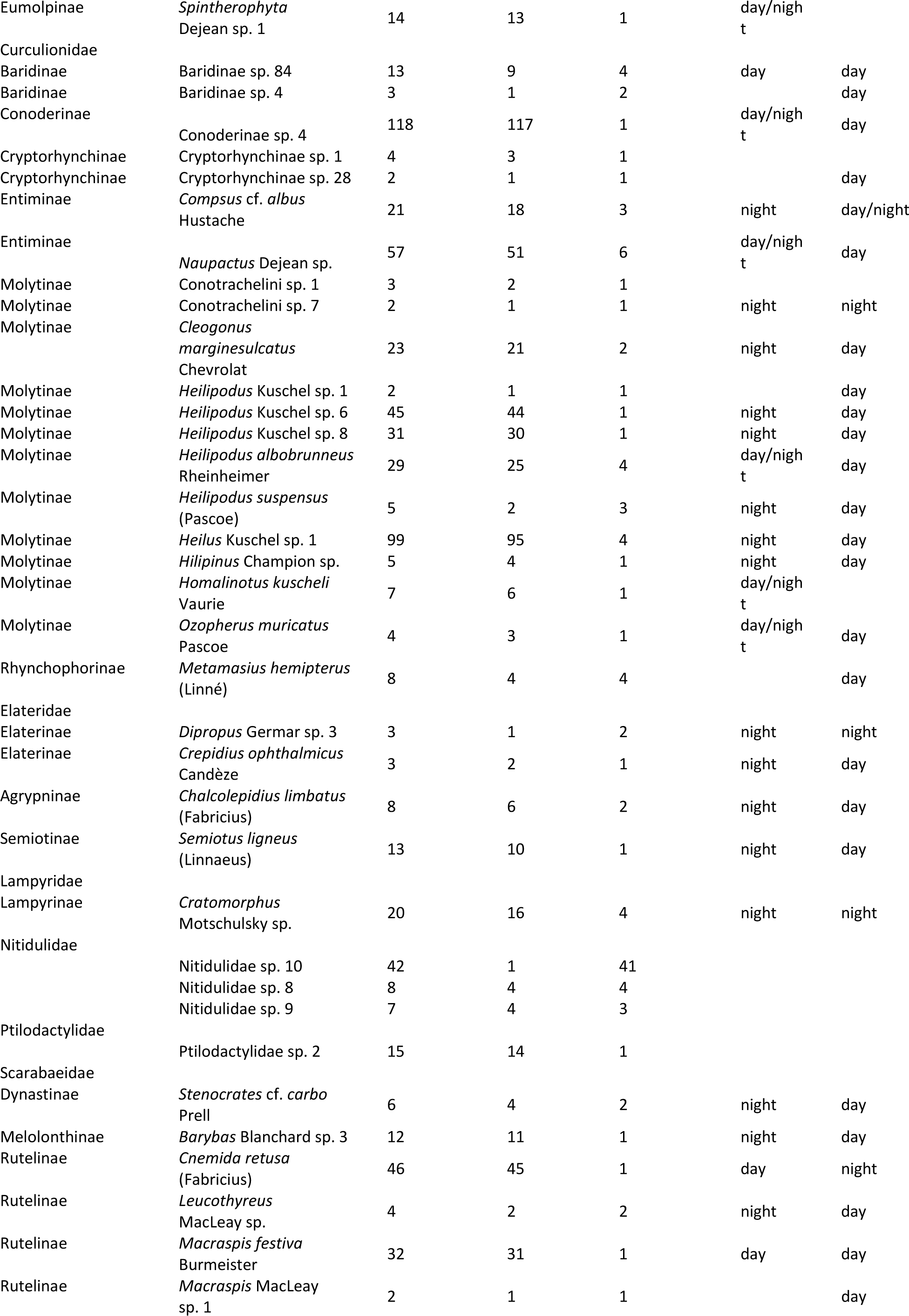
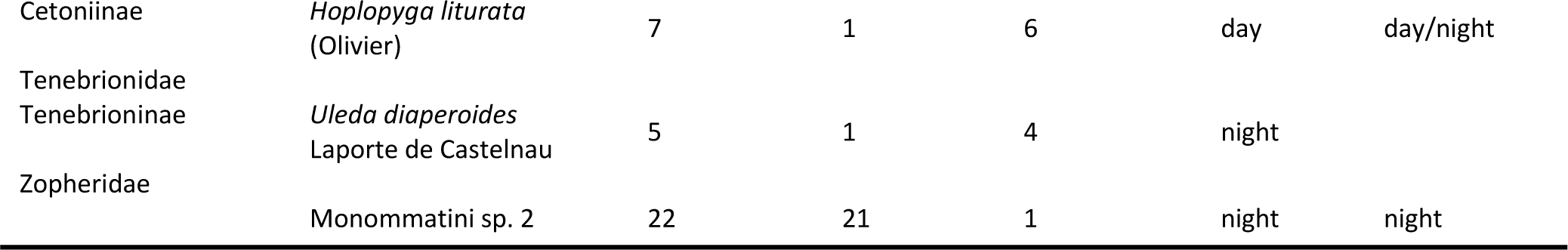
Adult beetle species collected on 23 canopy-tree species and in the understory in a lowland tropical rainforest in Venezuela, 1997–1999.

**Table 2.** Most species–rich beetle families (≥ 20 spp.) and beetles families collected on 23 canopy-tree species and in the understory in a lowland tropical rainforest in Venezuela, 1997–1999.

| Family | # canopy spp. | # shared spp. | % shared spp. | # spp. $n \geq 2$ | % shared spp. |
| --- | --- | --- | --- | --- | --- |
| Curculionidae | 248 | 20 | 8.1% | 143 | 14.0% |
| Chrysomelidae | 118 | 16 | 13.6% | 80 | 20% |
| Cerambycidae | 89 | 0 |  | 41 |  |
| Carabidae | 75 | 15 | 20.0% | 39 | 38.5% |
| Tenebrionidae | 37 | 1 | 2.7% | 16 | 6.3% |
| Scarabaeidae | 33 | 7 | 21.2% | 24 | 29.2% |
| Mordellidae | 22 | 0 |  | 18 |  |
| Nitidulidae | 22 | 3 | 13.6% | 12 | 25% |
| Elateridae | 20 | 4 | 20.0% | 11 | 36.4% |
| Buprestidae | 20 | 0 |  | 9 |  |
| Cantharidae | 15 | 1 | 6.7% | 10 | 10% |
| Zopheridae | 5 | 1 | 20% | 5 | 20% |
| Ptilodactylidae | 4 | 1 | 25% | 3 | 33.3% |
| Lampyridae | 4 | 1 | 25% | 1 | 100% |

**Table 3.** Beetle species of genera (≥ 2 spp.) collected on 23 canopy-tree species and in the understory in a lowland tropical rainforest in Venezuela, 1997–1999.

| Family | Subfamily | Genus | # total spp. | # canopy spp. | # understory spp. | # shared spp. |
| --- | --- | --- | --- | --- | --- | --- |
| Cantharidae | Silinae | *Ditemnus* LeConte | 3 | 3 | 1 | 1 |
| Carabidae | Haralinae | *Agra* Fabricius | 31 | 31 | 1 | 1 |
| Carabidae | Haralinae | *Apenes* LeConte | 4 | 2 | 4 | 2 |
| Carabidae | Haralinae | *Lebia* Latreille | 7 | 7 | 1 | 1 |
| Carabidae | Haralinae | *Loxandrus* LeConte | 3 | 2 | 3 | 2 |
| Carabidae | Haralinae | *Notiobia* Perty | 5 | 2 | 5 | 2 |
| Carabidae | Haralinae | *Pentagonica* Schmidt-Goebel | 2 | 2 | 1 | 1 |
| Carabidae | Haralinae | *Selenophorus* Dejean | 2 | 1 | 2 | 1 |
| Carabidae | Scaritinae | *Ardistomis* Putzeys | 2 | 2 | 1 | 1 |
| Carabidae | Scaritinae | Clivinina Rafinesque | 7 | 3 | 6 | 2 |
| Carabidae | Scaritinae | *Stratiotes* Putzeys | 2 | 1 | 2 | 1 |
| Chrysomelidae | Alticini | *Hypolampsis* Clark | 4 | 4 | 1 | 1 |
| Chrysomelidae | Alticini | *Omophoita* Chevrolat | 3 | 3 | 1 | 1 |
| Chrysomelidae | Galerucinae | *Diabrotica* Chevrolat | 16 | 16 | 6 | 6 |
| Chrysomelidae | Galerucinae | *Zepherina* Bechyné | 3 | 3 | 1 | 1 |
| Chrysomelidae | Eumolpinae | *Antitypona* Weise | 3 | 3 | 1 | 1 |
| Chrysomelidae | Eumolpinae | *Colaspis* Fabricius | 2 | 2 | 2 | 2 |
| Chrysomelidae | Eumolpinae | *Entomochirus* Lefèvre | 2 | 2 | 1 | 1 |
| Chrysomelidae | Eumolpinae | *Spintherophyta* Dejean | 3 | 3 | 1 | 1 |
| Curculionidae | Molytinae | *Heilipodus* Kuschel | 9 | 8 | 6 | 5 |
| Curculionidae | Molytinae | *Heilus* Kuschel | 5 | 5 | 1 | 1 |
| Curculionidae | Molytinae | *Homalinotus* Sahlberg | 3 | 3 | 1 | 1 |
| Elateridae | Elaterinae | *Crepidius* Candèze | 2 | 2 | 1 | 1 |
| Elateridae | Elaterinae | *Dipropus* Germar | 3 | 3 | 1 | 1 |
| Lampyridae | Lampyrinae | *Cratomorphus* Motschulsky | 2 | 2 | 1 | 1 |
| Scarabaeidae | Dynastinae | *Stenocrates* Burmeister | 4 | 3 | 2 | 1 |
| Scarabaeidae | Melolonthinae | *Barybas* Blanchard | 3 | 3 | 1 | 1 |
| Scarabaeidae | Rutelinae | *Cnemida* Kirby | 2 | 2 | 1 | 1 |
| Scarabaeidae | Rutelinae | *Macraspis* MacLeay | 8 | 7 | 3 | 2 |
| **TOTAL spp.** |  |  | **145** | **130** | **59** | **44** |

Therefore, the total number of species per genus of Chrysomelidae and Curculionidae might be larger than given in Table 3.

To analyze the statistical significance of the distribution of the taxa between the canopy and ground sample, I chose a two-sample paired test. I used the non-parametric Wilcoxon signed rank test due to non-normal distribution in PAST (Version 3.21; Hammer et al. 2001). After removing all rows with zero difference, the absolute values of the differences are ranked. The resulting W is the sum of the rank.

## Results

In total, 862 adult beetle (morpho)-species in 45 families were collected on 23 canopy-tree species during the one year of study. Of these, 70 species (8.1%) in 11 families were found also in the herb layer and/or on the ground (Table 1). Excluding all unique singletons out of the sample (n = 480), a proportion of 14.6% of all species was shared between canopy and understory. The majority of the 70 species, 61.4% (n = 43), was sampled with more individuals in the canopy, while 16 species (22.9%) were collected with more individuals in the understory (W = 1357.5, p < 0.001). Eleven species (15.7%) were represented by an equal proportion of canopy and understory individuals in the sample. Seventeen (24.3%) out of the 70 shared species were represented only as single individual in the canopy sample. In contrast, 38 shared species (54.3%) were collected only as single specimen in the understory.

Most species represented in the understory sample were Curculionidae (n = 20), Chrysomelidae (n = 16), and Carabidae (n =15) followed by Scarabaeidae (n = 7), Elateridae (n = 4), and Nitidulidae (n = 3) (Table 1). The families Cantharidae, Lampyridae, Ptilodactylidae, Tenebrionidae, and Zopheridae comprise only each one species represented in both strata. Among the 10 most species-rich beetle families with at least 20 species collected in the canopy of the 23 target tree species, Buprestidae, Cerambycidae, and Mordellidae were not found in the understory (Table 2). According to the number of species collected in the canopy, Scarabaeidae (21.2%), Carabidae (20%), and Elateridae (20%) have the highest proportion of shared species. Excluding all singletons (Table 2), 38.5% of Carabidae, 36% of Elateridae, and 29.2% of Scarabaeidae were shared between both strata.

Among Curculionidae, most shared species (n = 14 (70%); W = 132, p < 0.001) were collected with more individuals in the canopy (Table 1). The same applies to shared species of Chrysomelidae (n = 11 (68.75%); W = 79, p < 0.01). In contrast, most shared carabid species (n = 8 (53.3%); W = 72, p < 0.5) were represented with more individuals in the understory. Forty-four out of the 70 shared species belong to 29 identified genera sampled with at least two species in the crane plot (Table 3). These 29 genera in seven families comprise altogether 145 species with 130 species (89.7%) collected in the canopy and 59 species collected in the understory (W = 330, p < 0.01). Six out of 10 carabid genera (60%; W = 72, p < 0.5) were collected with more species in the understory. Except for the eumolpine genus *Colaspis* Fabricius, the genera of all other beetle families had more representatives in the canopy compared to the understory.

Some species were observed active in different strata of the forest (Table 1). Twenty-two species (31.4%) showed different activity in the canopy and understory, while seven species were active during the same period of the day in both strata. Of the 22 species with different activity in the canopy and understory, 21 species were active in the canopy during daytime. For instance, the ruteline species *Cnemida retusa* (Fabricius) diurnal in the canopy was found resting on the forest floor at night. In Elateridae, the nocturnal *Chalcolepidius limbatus* (Fabricius), *Semiotus ligneus* (Linnaeus), and *Crepidius ophthalmicus* Candèze regularly observed in the canopy during the night, were found on the ground, on the base of trunks or on herbs in the daytime. In Curculionidae, some nocturnal Molytinae, which were found only during the night in the canopy, were collected on the forest floor and understory, respectively, during the day (e.g., *Hilipinus* Champion sp., *Cleogonus marginesculatus* Chevrolat, *Heilus* Kuschel sp. 1, and three species of *Heilipodus* Kuschel). Among the galerucine genus *Diabrotica* Chevrolat, specimens of five nocturnal canopy species were found resting on herbs during the day.

## Discussion

### Sampling completeness

The data set comprises only a small proportion of the beetle diversity in the study area. The sampled 23 canopy-tree species represent less than one percent of the 3189 tree species known from the Venezuelan Amazon (Ter Steege et al. 2016). Moreover, canopy trap and hand collection were defined to only a small area of the outer tree crowns. Adult beetles were sampled from leaves, smaller twigs, fruits, and flowers neglecting stems or epiphytes on tree crowns. These are data from the two uppermost out of five commonly recognized strata in tropical rainforests (Richards 1971; Basset et al. 2015). In contrast, the understory is often defined as the vegetation layer between the tree canopy and the ground cover (Lawrence 2008) and can exceed a height of 10 m (Richards 1971). The beetle sample in the understory includes the soil surface / litter layer and the herb layer up to height of about 1.5 m. Although I sampled also on fruit falls of some monitored canopy trees, the plant composition and resource availability in the understory was different from those in the canopy. Still, I compare the strata use of adult beetles between the two ends of the vertical gradient with a distance of about 20 m. In addition, the data set comprises a full year of study. Thus, the sample should be representative for the beetle fauna of the Venezuelan study plot.

I combined active and passive methods in both strata, so that the methods complement each other and increase the chances of success of the inventory (Hyvarinen et al. 2006; Russo et al. 2011; Lamarre et al. 2012; Riley Peterson et al. 2021). Hand collecting is a good addition to the trap samples particularly for walking and unconcealed beetles. However, small individuals or cryptic species may not be noticed and other species may escape (Lopes et al. 2019). Nevertheless, canopy focus was adult beetle host associations comprising leaves, flowers, fruits, whereas understory samples included the ground, trunks and herbs. Similarly, the (semi-)quantitative trap collections focused on different guilds. The window traps installed in the tree crowns are particularly suitable for collecting small and active flying insects and thus, for canopy-inhabiting beetles of tropical rainforests that are usually good flyers (Basset et al. 1997; Lamarre et al. 2012). Otherwise, walking beetles are not captured with the flight interception traps. In contrast, pitfall trapping focuses on epigeal invertebrates, which are highly active and walking (Hoekman et al. 2017). Particularly predatory Carabidae and Staphylinidae are well represented in these traps (Barber 1931; Desender & Maelfait 1986; Honêk 1988; Jarosík 1992; Mommertz et al. 1996). Summarizing, sampling regimes in the understory and canopy focused largely on different guilds probably limiting the number of shared beetle species.

### General stratification

As a result, most beetle species in the sample (85.5%; excluding unique singletons) were found exclusively in the canopy. This in concordance with a study in a lowland rainforest in Cameroon, where arthropod densities and species richness were higher in the canopy than within the shrub layer (Basset et al. 1992). Only 70 out of 480 species (n ≥ 2) were collected at both canopy and understory level within the crane plot.

Microhabitat specialization is one reason for clear stratification of several taxa. For instance, most of the taxa of Curculionidae and Staphylinidae found in leaf litter were restricted to this habitat (Anderson & Ashe 2000). Hammond et al. (1997) identified 77 out of 140 weevil species as canopy specialists in a Sulawesi lowland rainforest.

Moreover, for most weevil species in Panama the vertical distribution in the forest was consistent from year to year (Wolda et al. 1998). Medianero et al. (2003) found among a leaf-mining guild only two out of 137 species and among gall makers only one out of 109 common to both canopy and understory. The proportion of gallers or miners in both the canopy and understory amounted to only 6% in this Panamanian wet forest. Climate conditions can largely contribute to high specificity of insect populations to distinct vertical strata within the forest. Vertical microclimatic and biotic gradients are much steeper in tropical rainforests than in temperate forests (Parker 1995; Hallé 1998) such as vertical complexity and stratification are better developed in tropical than in temperate forests (Smith 1973; Terborgh 1985). In addition to distinct microclimatic conditions, canopy and understory have different features in forest structure and resource availability (Grimbacher & Stork 2007; Sobek et al. 2009). Thus, stratification should be more pronounced in complex tropical forests.

Still, saproxylic beetles in temperate forests also show a clear vertical stratification (Vodka & Cizek 2013; Plewa et al. 2017). Ulyshen and Hanula (2007) collected with flight interception traps about 29% and 31% of species exclusively in the canopy or near the ground, respectively, remaining 40% shared between the two strata in a temperate deciduous forest in the USA. In an Italian temperate forest, 46 out of a total of 88 species were shared between canopy and ground layer, whereas 30% were caught only in the canopy and 30% exclusively in the ground layer (Hardersen et al. 2014). Otherwise, some studies located only few canopy specialists in tropical forests. Stork and Grimbacher (2006), for instance, found 72% of the species (excluding rare species: singletons and doubletons) in both strata whereby 24 and 27% of the abundant species were specialized to the canopy and the ground strata, respectively. One study in an Indonesian tropical forest revealed that only 8–13% of beetles are canopy specialists (Hammond et al. 1997). In Sulawesi, 56% of more abundant dung beetle species did not exhibit stratum preference, while 39% were specific to the canopy and 5% to the ground layer (Davis et al. 2011). Leksono et al. (2005) showed that few attelabid species clearly showed strong preference to specific layers and that the number of stratum specialist beetles was lower than generalist beetles. The proportion of shared species vs. stratum specialists just might strongly depend on the methods used to collect beetles, the exact vertical position of the traps in the forest and finally the structure of the forests. In this context, McCaig et al. (2019) suggested that the best fit for vertical stratification, is either the distance from ground or the distance down from the canopy.

### Specific strata use in guilds and families

Stratum specialization may apply to some taxa and guilds, respectively. In the Venezuelan sample, there are no shared species in Buprestidae, Cerambycidae, and Mordellidae, although these families were sampled with at least 20 species in the canopy. All Mordellidae and most species of Cerambycidae were collected exclusively on flowering trees (Kirmse & Chaboo 2020). These resources were not sampled in the understory (not available there) lowering the chance to collect these beetle families there. The same applies to Buprestidae, which were either flower visitors or leaf feeders in the canopy (Kirmse unpub. data). Although, there is a variety of larval feeding habits in these three families, most larvae feed internally in stems and leaves of plants or in dead wood (Liljeblad 1945; Bellamy & Nelson 2002; Bezark & Monné 2013). Such larval resources are largely available in the canopy. Thus, Buprestidae, Cerambycidae, and Mordellidae could complete their entire life cycle in the canopy. In contrast, only about 30% of the saproxylic beetle species were recorded only at the canopy stratum in temperate forests (Bouget et al. 2011). However, strata use might be associated with the season. Berkov and Tavakilian (1999) found that the xylophagous Cerambycidae *Palame* Bates spp. utilizing Lecythidaceae trees in French Guiana reproduced exclusively in the canopy in the wet season with reproduction at ground level restricted to the dry season.

The species-rich phytophagous beetle families Chrysomelidae, Curculionidae, Elateridae, and Scarabaeidae were represented as well with many species on flowering canopy trees (Kirmse & Chaboo 2020), but their dietary associations with the canopy trees were broader (Kirmse & Chaboo 2018; Kirmse & Ratcliffe 2019; Kirmse & Johnson 2020). These four families comprise a proportion of 14–36% of species shared between both sampled strata in the crane plot. The percentage of 14% in Curculionidae is lower in the Venezuelan sample compared to a study in Panama, where no more than 20% of curculionids were restricted to either canopy or ground stratum (Wolda et al. 1998). Yet, the 20% shared leaf-beetle species are in the range of the proportion of chrysomelid species in Panama, where 16% and 28%, respectively, were shared between the canopy and understory of wet and dry sites (Charles & Basset 2005). This Panama sample included 33–53% of species unique to the canopy.

In Chrysomelidae, Elateridae, and Scarabaeidae there are high proportions of larvae developing in the soil. Particularly most larvae of Eumolpinae and Galerucinae (Chrysomelidae) live in the soil feeding on roots in tropical rainforests (Jolivet 1994). Pokon et al. (2005) reared a total of 100 species of root-feeding chrysomelid larvae from the roots in a secondary tropical forest in New Guinea. Almost 90% of adults in the forest canopy were recruited from the species with root-feeding larvae. In the Venezuelan study, shared species of Chrysomelidae belong to Eumolpinae and Galerucinae including Alticini. In contrast, there were no shared species in the subfamily Cryptocephalinae, although represented with the same high number of species in the canopy such as Eumolpinae (Kirmse & Chaboo 2018). Many larvae of melolonthines, rutelines, and dynastines (Scarabaeidae) feed on roots (Ritcher 1958; Scholz & Chown 1995) as well as some Elateridae (Johnson 1999). In Curculionidae, larvae of the subfamily Entiminae feed externally in the soil on roots, while most larvae of Cryptorhynchinae and Molytinae feed in dead wood or other dead and decaying plant material (Anderson 2002). Other weevil groups, such as several Conoderinae emerged often from branches at canopy stratum (Berkov 2018). In the Venezuelan sample, most shared species belong to Molytinae. Generally, oviposition on the forest floor and hatching of larvae from the soil will enhance the probability to collect these taxa at or near the ground.

The occurrence of some shared beetle species coincided with resource availability on the forest floor. The shared species of Nitidulidae were collected with the pitfall traps on fruit falls. The same applies to the darkling beetle *Uleda diaperoides* Laporte de Castelnau and the zopherid Monommatini sp. 2. Carabidae revealed the highest proportion of shared species in the Venezuelan sample. Most shared species were collected with the pitfall traps on the fruit falls including the seed-feeding species of *Notiobia* Perty and *Selenophorus* Dejean (Arndt & Kirmse 2002). As many adults and larvae of ground beetles live in soil, in leaf litter or are active on the ground surface (Arndt et al. 2005), there is a good chance to collect most species with pitfall traps. In addition, many carabid species should be able to forage in different strata of the forest as adults of most species are omnivorous (Thiele 1977), even though carnivorous nutrition seems to prevail with 73.5% (Larochelle 1990). Taking into account that I sampled particularly ground beetles probably quite thoroughly with the pitfall traps, I will be able to evaluate which genera contain more arboreal than ground dwelling species. While the lebiine genera *Agra* Fabricius and *Lebia* Latreille were collected with most species only in the canopy, most other harpaline and scaritine genera revealed more species inhabiting the forest floor compared to the canopy indicating their strong association with the ground layer.

### Strata use and stratum switch

Stratum preferences of adult beetles might not only depend on the availability of food resources and larval substrates, but also on their diel activity and the microclimatic conditions in the forest layers. Twenty-two shared species showed distinct different activity in the canopy and understory, respectively. The use of different strata according to phases of activity is described by Erwin (2013). He found that Alleculinae feed on lichens and moss on tree trunks at night and spend the day hiding in suspended dry leaves elsewhere in the forest. Erwin (2013) suggested that many adult beetle species found in the forest canopy during the day hide and rest at night in the understory. Indeed, detailed studies in the canopy plot revealed that extrafloral nectar- and flower-visiting beetles visited their host trees only during a distinct period of the day (Kirmse & Chaboo 2019, 2020) causing a permanent diel migration. If there is a high proportion of beetle species changing their used stratum in the course of the day, the proportions of stratum generalists should increase if beetles are collected day and night. This might be supported by Basset et al. (2015), who found that the vertical turnover of arthropods particularly in phytophages and predators exceeds the horizontal and seasonal turnover in the rainforest in Panama.

One reason for the movement between strata according to phases of activity may be that there are most probably special adaptations required particularly for diurnal canopy specialists. Tree canopies are exposed habitats, and during the day they can be hot, dry, and receive high solar insolation (Parker 1995; Hallé 1998; Nakamura et al. 2017; Frenne et al. 2019). As a result, fluctuations in relative humidity and air temperature, and wind speed are noticeably higher in the upper canopy than in the understory (Pinker 1980; Parker 1995; Walsh 1996; Szarzynski & Anhuf 2001; Turton & Siegenthaler 2004; Ulyshen 2011). Additionally, water condensation at night is frequent within the upper canopy, but absent in the understory (Blanc 1990). During the night, heat is lost through radiation and sensible as well as latent heat via condensation. Accordingly, temperature fluctuated in the Surumoni crane plot by almost 10°C in the upper canopy and only slightly near the ground (Anhuf & Rollenbeck 2001; Szarzynski & Anhuf 2001). Furthermore, light intensities can be 500” higher in the canopy of tropical moist forests than in the understory (Kelber et al. 2006). Strategies to overcome the hygrothermal stress during the day and water condensation during the night in the canopy are feeding at night in the upper canopy and resting during the day in lower layers. Indeed, 21 out of 22 beetle species with different activity between both strata were observed active in the canopy during the night. Basset et al. (2003a), for instance, found Curculionidae more active during the day in the upper canopy and faunal turnover in the upper canopy between day and night was higher compared to the understory. These climatic conditions might largely contribute to a steady interchange of the canopy fauna.

## Conclusions

The proportion of shared beetle species between the canopy and the understory was with 14.6% low. Beetle families with many species represented in both strata contain many species with larvae developing in the soil: Chrysomelidae, Curculionidae, Elateridae, and Scarabaeidae. Carabidae as commonly omnivore or predatory group revealed the highest proportion of shared species. Moreover, among the beetle families sampled with at least 20 species on the 23 canopy trees, most carabid genera contained more species sampled in the understory compared to the canopy. In contrast, the mainly flower-visiting beetle families Cerambycidae and Mordellidae as well as the Buprestidae were collected exclusively in the canopy. Their utilized resources were not sampled/available in the understory. In addition, much of the larval feeding habits in these three families occur internally in living plant parts or in dead wood, which is available in the canopy. Thus, much of the strata use found in the Surumoni crane plot sample can be attributed to resource availability and larval development.

## Acknowledgements

For the determination of the beetles I thank all the beetle experts indicated in the Methods. I cordially thank Jens Wesenberg for superior assistance in all botanical matters and field work. The Austrian Academy of Sciences and colleagues are gratefully acknowledged for their support and permission to join the Surumoni project in Venezuela. The fieldwork was supported by grants from the ESF Tropical Canopy Programme and the Stiftung der Deutschen Wirtschaft, Germany. Finally, but most important, the Leopoldina, Germany, supported generously this study by scientific and financial means.

